# Contemporary seasonal human coronaviruses display differences in cellular tropism compared to laboratory-adapted reference strains

**DOI:** 10.1101/2025.04.13.648626

**Authors:** Matthew J Gartner, Monique L Smith, Clyde Dapat, Yi Wen Liaw, Thomas Tran, Randy Suryadinata, Joseph Chen, Guizhi Sun, Rory A Shepherd, George Taiaroa, Michael Roche, Wen Shi Lee, Philip Robinson, Jose M Polo, Kanta Subbarao, Jessica A Neil

## Abstract

Seasonal human coronaviruses (sHCoVs) cause 15-30% of common colds. The reference strains used for research were isolated decades ago and have been passaged extensively but contemporary sHCoVs have been challenging to study as they are notoriously difficult to grow in standard immortalized cell lines. Here we addressed these issues by utilizing primary human nasal epithelial cells (HNECs) and immortalized human bronchial epithelial cells (BCi) differentiated at an air-liquid interface, as well as human embryonic stem cell-derived alveolar type II (AT2) cells to recover contemporary sHCoVs from human nasopharyngeal specimens. From 21 specimens we recovered four 229E, three NL63 and eight OC43 viruses. All contemporary sHCoVs showed sequence differences from lab-adapted CoVs, particularly within the spike gene. Evidence of nucleotide changes in the receptor binding domains within 229E and detection of recombination for both 229E and OC43 isolates was also observed. Importantly, we developed methods for the amplification of high titre stocks of NL63 and 229E, that maintained sequence identity, and we established methods for the titration of contemporary sHCoV isolates. Comparison of lab-adapted and contemporary strains in immortalised cell lines and airway epithelial cells revealed differences in cell tropism, growth kinetics and cytokine production between lab-adapted and contemporary sHCoV strains. These data confirm that contemporary sHCoVs differ from lab-adapted reference strains and, using the methods established here, should be used for study of CoV biology and evaluation of medical countermeasures.

**Importance:** Zoonotic coronaviruses have caused significant public health emergencies. The occurrence of a similar spillover event in the future is likely and efforts to further understand coronavirus biology should be a high priority. Several seasonal coronaviruses circulate within the human population. Efforts to study these viruses have been limited to reference strains isolated decades ago due to the difficulty in isolating clinical isolates. Here, we use human airway and alveolar epithelial cultures to recover contemporary isolates of NL63, 229E and OC43. We establish methods to make high titre stocks and titrate 229E and NL63 isolates. We show that contemporary isolates of NL63 and OC43 have a different tropism within the respiratory epithelium compared to lab-adapted strains. Although 229E clinical and lab-adapted strains similarly infect the respiratory epithelium, differences in host response and replication kinetics are observed. Using the methods developed here, future research should include contemporary isolates when studying coronavirus biology.

## Introduction

Coronaviruses (CoVs) are ubiquitous in a wide range of avian species and mammals, which can act as a reservoir for transmission to humans (1). Seven coronaviruses are known to infect humans via the respiratory tract causing a spectrum of respiratory disease. In the past two decades, three zoonotic CoVs, severe acute respiratory syndrome coronavirus (SARS-CoV), middle east respiratory syndrome coronavirus (MERS-CoV) and SARS-CoV-2 have emerged from animals to cause significant public health emergencies (2, 3). In addition, four seasonal human CoVs (sHCoVs; 229E, NL63, OC43 and HKU1) have been circulating in the human population for decades and are presumed to have zoonotic origins (4). sHCoVs generally cause self-limiting respiratory infections and account for 15-30% of common cold cases (5). However, these viruses can lead to more severe respiratory disease in neonates, the elderly and immunocompromised populations (6, 7).

While there has been a significant research effort to characterise and understand the recently emerging CoVs (SARS-CoV, MERS-CoV and SARS-CoV-2), we have less knowledge of the biology and evolutionary dynamics of the sHCoVs. Like emerging CoVs, sHCoVs display genetic evolution over time predominantly within the Spike (S) protein, suggesting emergence of viral variants with altered tropism or the ability to escape pre-existing antibody immunity, may occur (8–10). Studies investigating host cell tropism, entry, replication and assessment of medical countermeasures using sHCoVs are limited primarily to lab-adapted reference strains. These lab-adapted reference strains were isolated decades ago and have been extensively passaged in immortalised cell lines. Whether these lab-adapted strains accurately represent the contemporary strains circulating in the population is unclear. Isolation and propagation of contemporary sHCoVs from clinical specimens in immortalized cell lines has been challenging. A previous study published by Dijkman et al in 2013 reported the use of human bronchial airway epithelial cells differentiated at an air-liquid interface (ALI) to isolate 10 contemporary sHCoV strains (11). This study demonstrated that one universal cell culture system could be used to isolate diverse sHCoV strains. However, the success rate of recovery of NL63 isolates was lower than the other viruses, likely because the requirements for growth of each sHCoV differs. There was also limited characterisation of sequence variability and replication of the contemporary isolates relative to lab-adapted strains.

Therefore, to overcome these issues and establish models to study contemporary sHCoVs, we have built on the methods of Dijkman et al to use a combination of three models of the human respiratory tract; primary human nasal epithelial cells (HNECs) differentiated at an ALI, immortalised human bronchial epithelial cells (BCi) differentiated at an ALI, and embryonic stem-cell derived alveolar type II cells (AT2), to recover novel contemporary sHCoV isolates. We leveraged this suite of respiratory tract models (12) to isolate 15 contemporary sHCoVs, including 229E (n=4), NL63 (n=3) and OC43 (n=8) from 21 nasopharyngeal swab specimens that were PCR-positive for sHCoV nucleic acids. We used next generation sequencing to generate full length genome sequences for each isolated sHCoV and performed phylogenetic analysis to understand genetic diversity among the isolates and published sequences. In addition, we established protocols for generating high titre virus stocks using immortalised cell lines and compared replication kinetics of the contemporary and laboratory-adapted reference sHCoV strains in cell lines and complex airway models.

## Results

### Isolation of contemporary sHCoVs using primary and stem-cell derived cell culture systems

We attempted to recover infectious virus from 21 nasopharyngeal specimens that were PCR+ for non-HKU1 sHCoVs (229E (n=5), OC43 or NL63 (n=16)) using human embryonic stem cell-derived AT2 cells, immortalised BCi cells differentiated at an ALI or primary HNECs differentiated at an ALI (Fig. 1A, Table 1). Nasopharyngeal specimens were collected over a period of 6 years, from 2017 to 2022. In brief, clinical specimens were diluted in phosphate buffered saline (PBS) before incubation with the cells for 2 hours at 33°C. Following removal of inoculum, cells were cultured at 33°C for up to 7 days and supernatant (apical wash for ALI cultures) collected at various timepoints post-infection. For most experiments, supernatants from replicate wells were pooled to generate a virus stock and assessed for the presence of sHCoV nucleic acids by qRT-PCR. Two samples were lost due to contamination (likely due to presence of yeast in the specimen). Of the remaining 19, four samples had no detectable levels of any virus at any timepoint when inoculated onto AT2 or BCi cells as determined by qRT-PCR (data not shown). Infectious sHCoVs were recovered from the remaining 15 specimens.

**Figure 1.**
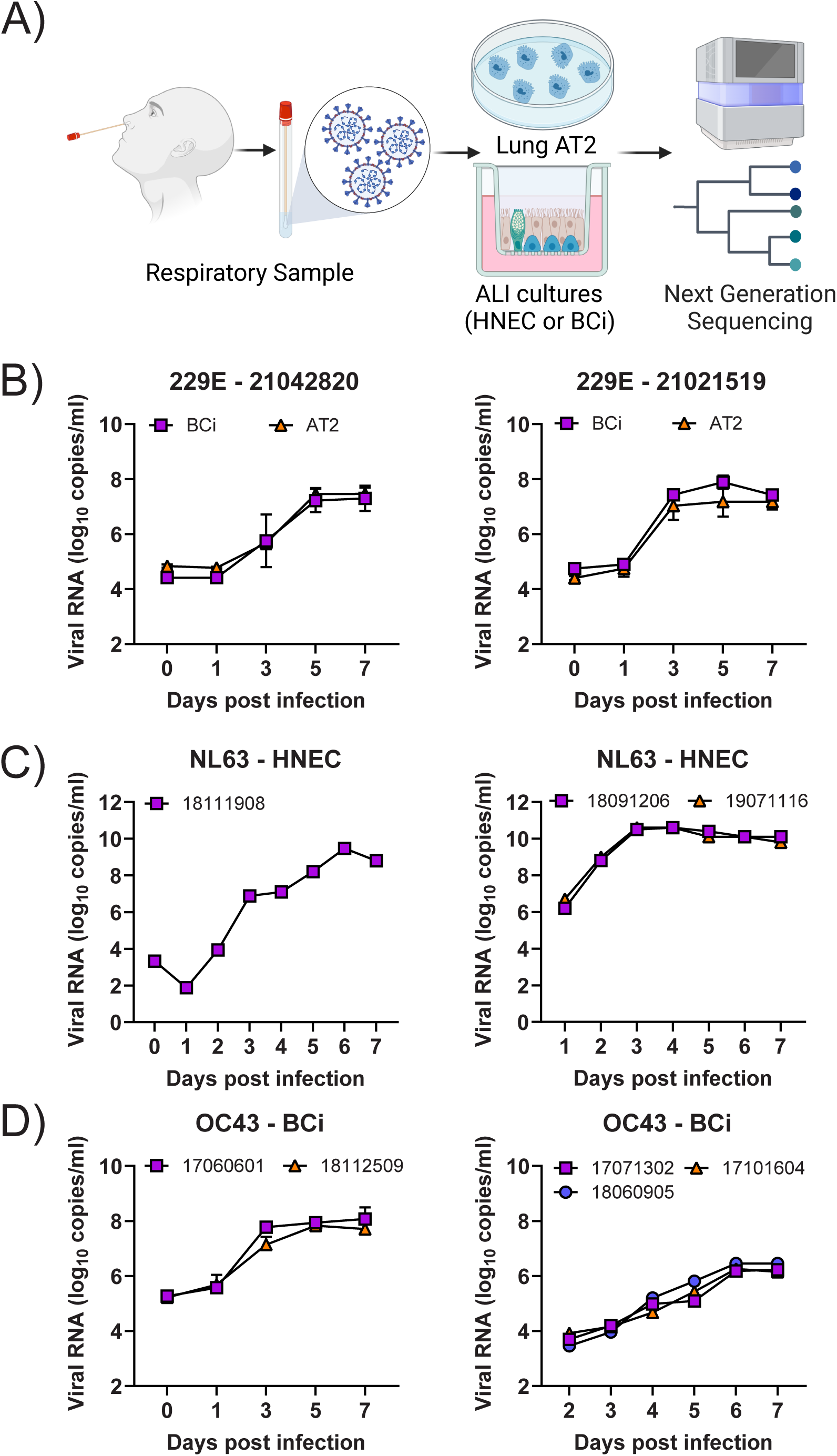
Isolation of contemporary sHCoVs. A) Schematic of the procedure for recovering contemporary sHCoVs from human nasal swabs. Replication kinetics (Log_10_ genome copies/mL from 0 to 7 dpi) of recovered 229E isolates in immortalised BCi cells grown at an ALI and H9-derived AT2 cells (B, C), NL63 isolates in primary HNECs grown at an ALI (D, E) and OC43 isolates in immortalised BCi cells grown at ALI (F, G). For (B), (C) and (F), the mean ± SD from 4 un-pooled replicates in shown. For (D), (E) and (G) the mean from 4 pooled replicates is shown. Each graph represents data generated from a single attempt to recover virus using a nasal swab specimen.

**Table 1:**
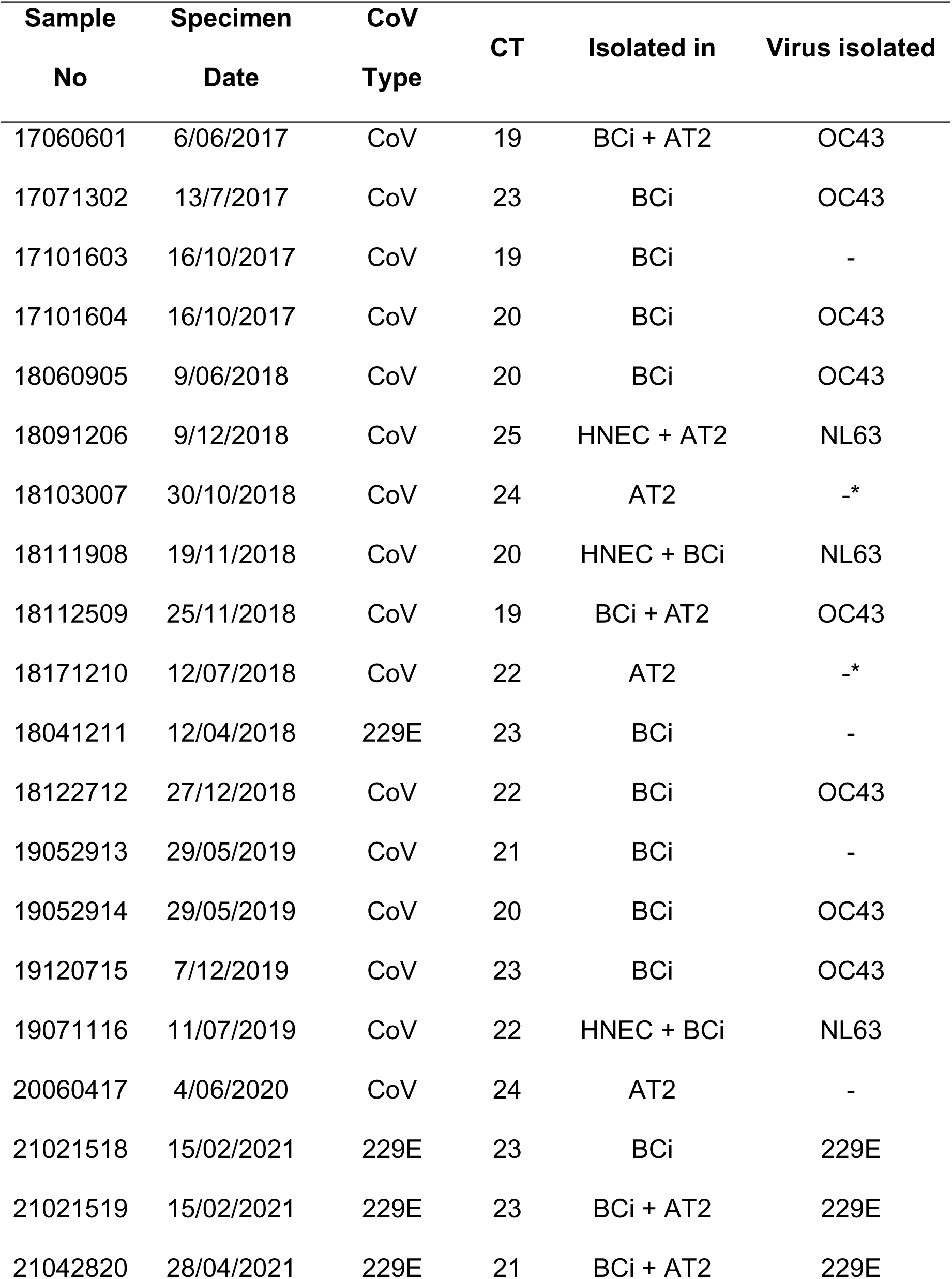
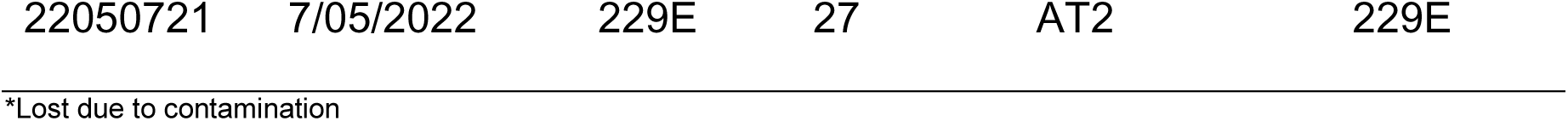
Clinical specimens for isolation of seasonal coronaviruses.

The five 229E+ specimens were inoculated on AT2 and/or BCi cells. As shown in Table 1, two samples were cultured in BCi cells only, one in AT2 cells only and two in both cell types. Of the samples cultured in BCi cells only, one sample collected in 2018 did not yield virus. Infectious 229E was recovered from all other samples collected in 2021 and 2022. Samples 21042820, 21021519 and 22050721 grew robustly in AT2 cells peaking at 5 to 7 days post-infection (dpi, Fig. 1B and data not shown). Cytopathic effect was observed from 3 dpi (data not shown). For samples 21021518, 21021519 and 21042820, similar growth kinetics were observed when virus was recovered in BCi cells (Fig. 1B and data not shown). No obvious CPE was observed in BCi cells that yielded virus (data not shown). Overall, 80% (4/5) of clinical samples containing 229E RNA yielded infectious virus from AT2 and BCi cells, with similar titres.

Three 229E negative specimens yielded NL63: samples 18091206 and 18111908 collected in 2018 and 19071116 collected in 2019. While initial attempts to recover virus from sample 18091206 in AT2 cells or 18111908 and 19071116 in BCi cells were unsuccessful, low levels of virus were detectable in either the inoculum or on 1 dpi in these cultures (Sup. Fig. 1A). Given that NL63 primarily causes an upper respiratory tract infection and infects ACE2 expressing ciliated cells, we re-attempted virus isolation from these clinical samples using HNECs. All three viruses grew in HNECs with virus peaking on 6 dpi for 18111908 and 3 dpi for 18091206 and 19071116 (Fig. 1C). The differential viral replication kinetics between 18111908 compared to 18091206 and 19071116 may be explained by different HNEC donors used for virus isolation. In our hands, recovery of contemporary NL63 viruses is most robust when using nasal epithelial cells compared to cells from lower in the respiratory tract.

The remaining 8 clinical samples yielded OC43. As shown in Table 1, they included three samples collected in 2017 (17060601, 17071302 and 17101604), three collected in 2018 (18060905, 18112509 and 18122712) and two collected in 2019 (19052914 and 19120715). Samples 17060601 and 18112509 were cultured in BCi and AT2 cells while the remaining were only cultured in BCi cells. Attempts to recover samples 17060601 and 18112509 in AT2 cells were unsuccessful (Sup. Fig. 1B). All viruses were recovered in BCi cells, with virus peaking at day 3-6 dpi (Fig. 1D). This outcome was the opposite to that observed for the lab-adapted OC43 strain (VR-1558) which showed robust replication in AT2 cells but no infection in BCi cells (Sup. Fig.1C-D). This suggests that clinical OC43 strains may show differences in host susceptibility and respiratory tract tropism compared to the lab-adapted reference strain.

### Genetic characterisation of contemporary sHCoVs

To study the evolutionary relationships between the contemporary sHCoV isolates and other circulating CoVs, we generated full genome sequences for each contemporary sHCoV and built phylogenetic trees including publicly available genomes. For 229E, two isolates (21021519 and 21021518) had identical genome sequences and identical sample collection dates, so we removed 21021518 from further analysis. Consistent with previous studies, the 229E tree had a ladder-like shape with short terminal branches and sequence divergence was proportional to virus isolation date, consistent with genetic drift (8–10, 13, 14). Our 229E isolates 21042820, 21021519 and 22050721 all fell within one clade (genotype 7a) containing sequences from 2017 to 2023 from diverse geographical locations, including Russia, Japan, Italy and Tanzania (Fig. 2A), consistent with previous reports showing that 229E sequences cluster by date and not location (8, 14, 15). We noted two distinct clades of sequences with collection dates ranging from 2016 to 2023 (denoted genotypes 7a and 7b), suggesting the co-circulation of two different genotypes of 229E.

**Figure 2.**
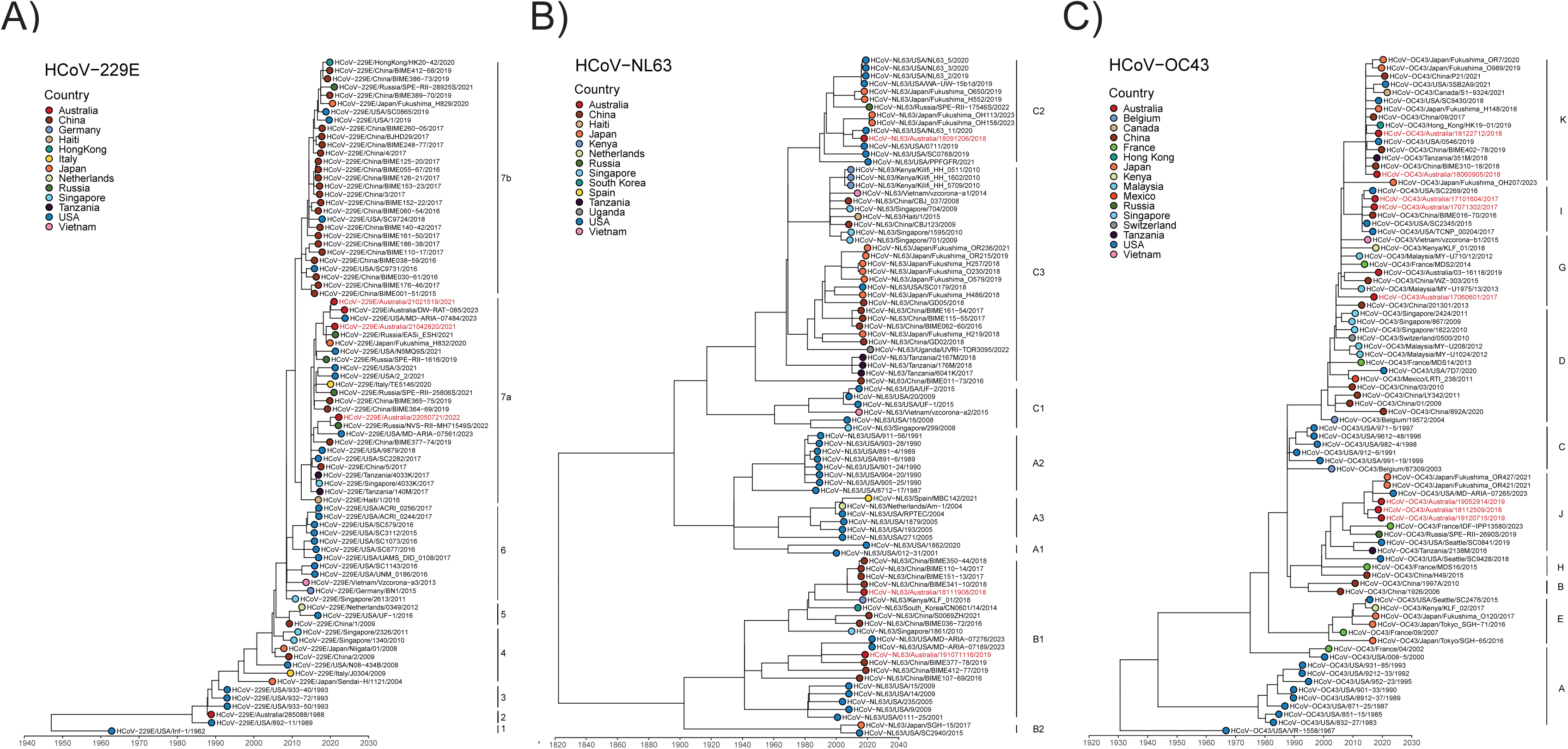
Genetic characterisation of sHCoVs. Time-resolved maximum likelihood phylogenetic analysis of full genome nucleotide sequences of contemporary 229E (A), NL63 (B) and OC43 (C) isolates compared to publicly available sequences. Sequences from viruses isolated in this study are shown in red text. Coloured symbols indicate different countries of origin.

Analysis of NL63 sequences showed our isolates grouped into two distinct genotypes; 18111908 and 19071116 clustered with sequences in genotype B1, while 18091206 grouped with sequences in genotype C2 (Fig. 2B). Similar to the 229E tree and consistent with other studies (9, 10, 13, 14, 16), the OC43 tree showed sequences that diverged based on isolation date and not location, suggestive of genetic drift. Our OC43 sequences showed the eight isolates grouped into four co-evolving lineages (Fig. 2C). Samples 18112509, 19052914 and 19120715 grouped with genotype J sequences, 17060601 grouped with genotype G, 17071302 and 17101604 grouped with genotype I and 18060905 and 18122712 grouped with genotype K sequences (Fig. 2C).

We performed Simplot analysis and calculated nucleotide identity between each isolate compared to the respective reference strain, to probe which viral genes demonstrated the greatest diversity. Simplot analysis of all three viruses demonstrated the highest genetic diversity within the S gene (Fig. 3A-C), with 229E isolates showing 95.51-95.80 percent identity within S, NL63 showing 96.05-98.16 percent identity within the S (Table 2) and OC43 showing 95.50-96.16 percent identity within the S compared to the respective lab-adapted reference strain (Table 3).

**Figure 3.**
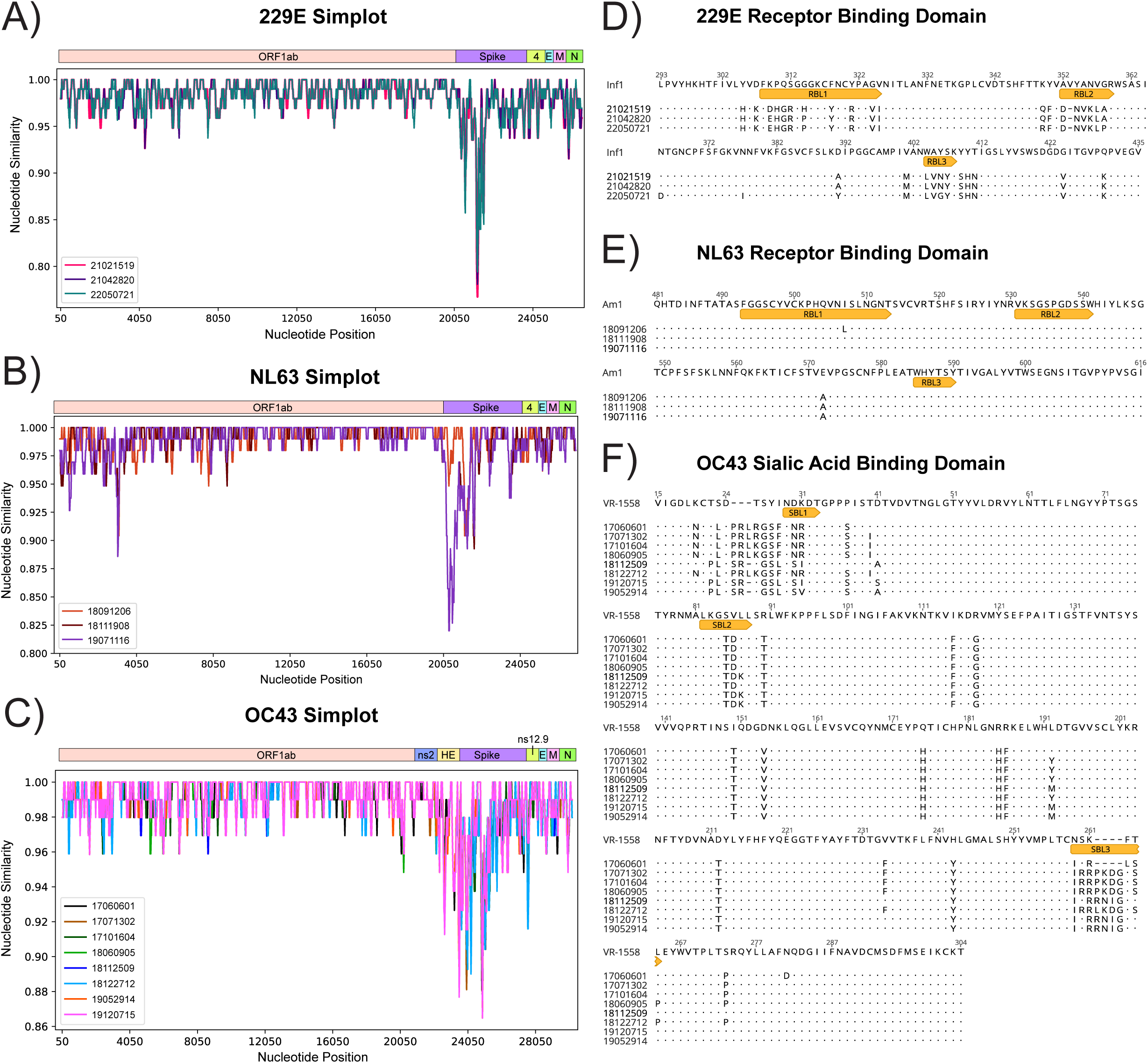
Sequence analysis of sHCoVs. SimPlot analysis of full genome sequences of contemporary 229E (A), NL63 (B) and OC43 (C) isolates showing the percent nucleotide similarity across nucleotide positions compared to reference strains (HCoV-229E/USA/Inf1/1962, HCoV-NL63/NED/Amsterdam1/2004 and HCoV-OC43/USA/VR-1558/1967, respectively). Analysis of the 229E receptor binding domain (D), NL63 receptor binding domain (E) and OC43 sialic acid binding domain (F) from contemporary isolates compared to the reference strain. RBL: Receptor Binding Loop, SBL: Sialic Acid Binding Loop.

**Table 2:**
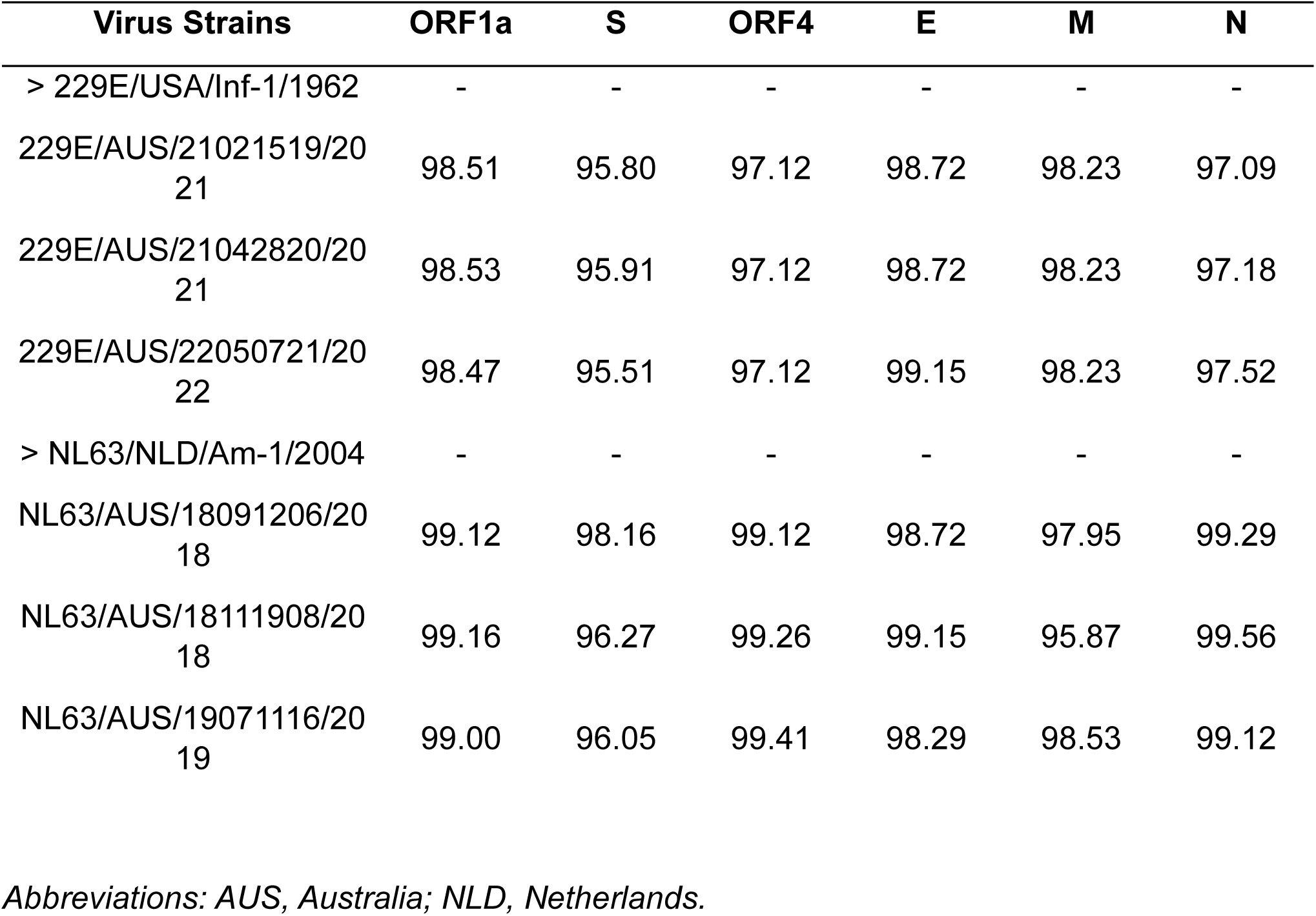
Nucleotide identity of clinical isolates compared to HCoV-229E and HCoV-NL63 reference strains.

| Virus Strains | ORF1a | S | ORF4 | E | M | N |
| --- | --- | --- | --- | --- | --- | --- |
| > 229E/USA/Inf-1/1962 | - | - | - | - | - | - |
| 229E/AUS/21021519/2021 | 98.51 | 95.80 | 97.12 | 98.72 | 98.23 | 97.09 |
| 229E/AUS/21042820/2021 | 98.53 | 95.91 | 97.12 | 98.72 | 98.23 | 97.18 |
| 229E/AUS/22050721/2022 | 98.47 | 95.51 | 97.12 | 99.15 | 98.23 | 97.52 |
| > NL63/NLD/Am-1/2004 | - | - | - | - | - | - |
| NL63/AUS/18091206/2018 | 99.12 | 98.16 | 99.12 | 98.72 | 97.95 | 99.29 |
| NL63/AUS/18111908/2018 | 99.16 | 96.27 | 99.26 | 99.15 | 95.87 | 99.56 |
| NL63/AUS/19071116/2019 | 99.00 | 96.05 | 99.41 | 98.29 | 98.53 | 99.12 |

*Abbreviations: AUS, Australia; NLD, Netherlands.*

**Table 3:** Nucleotide identity of clinical isolates compared to HCoV-OC43 reference strain VR-1558.

| Virus Strains | ORF1<br>ab | Ns2 | HE | S | Ns12.<br>9 | E | M | N |
| --- | --- | --- | --- | --- | --- | --- | --- | --- |
| ><br>OC43/USA/VR1558/19<br>67 | - | - | - | - | - | - | - | - |
| OC43/AUS/17060601/2<br>017 | 99.25 | 98.92 | 97.49 | 95.78 | 99.39 | 99.62 | 98.27 | 98.52 |
| OC43/AUS/17071302/2<br>017 | 99.23 | 98.92 | 97.33 | 95.50 | 99.09 | 99.62 | 98.27 | 98.52 |
| OC43/AUS/17101604/2<br>017 | 99.19 | 99.16 | 97.25 | 95.55 | 99.09 | 99.62 | 98.27 | 98.59 |
| OC43/AUS/18060905/2<br>018 | 99.21 | 99.16 | 97.18 | 95.47 | 99.09 | 99.62 | 98.27 | 98.66 |
| OC43/AUS/18112509/2<br>018 | 99.20 | 99.16 | 97.02 | 96.19 | 98.79 | 96.93 | 98.56 | 98.29 |
| OC43/AUS/18122712/2<br>019 | 99.20 | 99.16 | 97.10 | 95.50 | 99.09 | 99.23 | 98.27 | 98.74 |
| OC43/AUS/19052914/2<br>019 | 99.20 | 99.16 | 97.10 | 96.16 | 98.79 | 96.93 | 98.56 | 98.44 |
| OC43/AUS/19120715/2<br>019 | 99.18 | 99.16 | 96.94 | 96.13 | 98.79 | 96.93 | 98.56 | 98.37 |

*Abbreviations: AUS, Australia*

Given the diversity within S for 229E, NL63 and OC43, we generated alignments of predicted amino acid sequences for the S1 domain (Sup Fig. 2A-C) and the receptor binding domain (RBD) of 229E and NL63 (Fig. 3D-E) and the sialic acid binding domain (domain A) of OC43 (Fig. 3F), comparing our isolates to the appropriate reference strain. Analysis of the RBD for 229E isolates showed several amino acid changes within the receptor binding loops (RBLs) 1, 2 and 3 compared to reference strain, Inf1 (Fig. 3D). The three RBLs are responsible for binding to human amino peptidase N (hAPN) for entry and undergo substantial genetic variation (8, 17). Outside the RBD, we found several amino acid changes within the N-terminal domain (NTD) compared to the reference strain (Sup Fig. 2A), a region previously shown to be immunodominant in 229E S (18).

In contrast to 229E, we found minimal amino acid changes within the RBD of our NL63 isolates compared to the reference strain Am-1 (Fig. 3E). Instead, we found substantial variation within the N-terminal Unique domain of S1 (Sup Fig. 2B). Our OC43 isolates showed substantial variation compared to the reference VR1558 strain near sialic acid binding loop 1 (SBL1), with three of the eight isolates having a SR-TG sequence and the remaining five having a PRLRG insertion (Fig. 3F). In addition, all eight isolates showed variation in SBL2 and SBL3 compared to VR1558. The eight OC43 isolates also showed substantial variation within domain B compared to VR1558 (Sup Fig. 2C).

We constructed phylogenetic trees using S gene sequences for 229E, NL63 and OC43 to understand molecular evolution of the Spike (S) gene over time (Fig. 4A-C). Similar to the trees based on the whole genome, we found two co-circulating genotypes of 229E S sequences over the same timeframe (2016–2023), indicating two genotypes were circulating globally. Inconsistent with the full-length tree, 22050721 grouped with genotype 7b while 21021519 and 2106799 grouped with genotype 7a, suggesting 22050721 may be a recombinant. Recombination analysis using the sequences used to generate the phylogenetic tree predicted a recombination event in 22050721 between genotype 7a and 7b sequences with breakpoints occurring between ORF1ab and S (Sup Fig. 3A).

**Figure 4.**
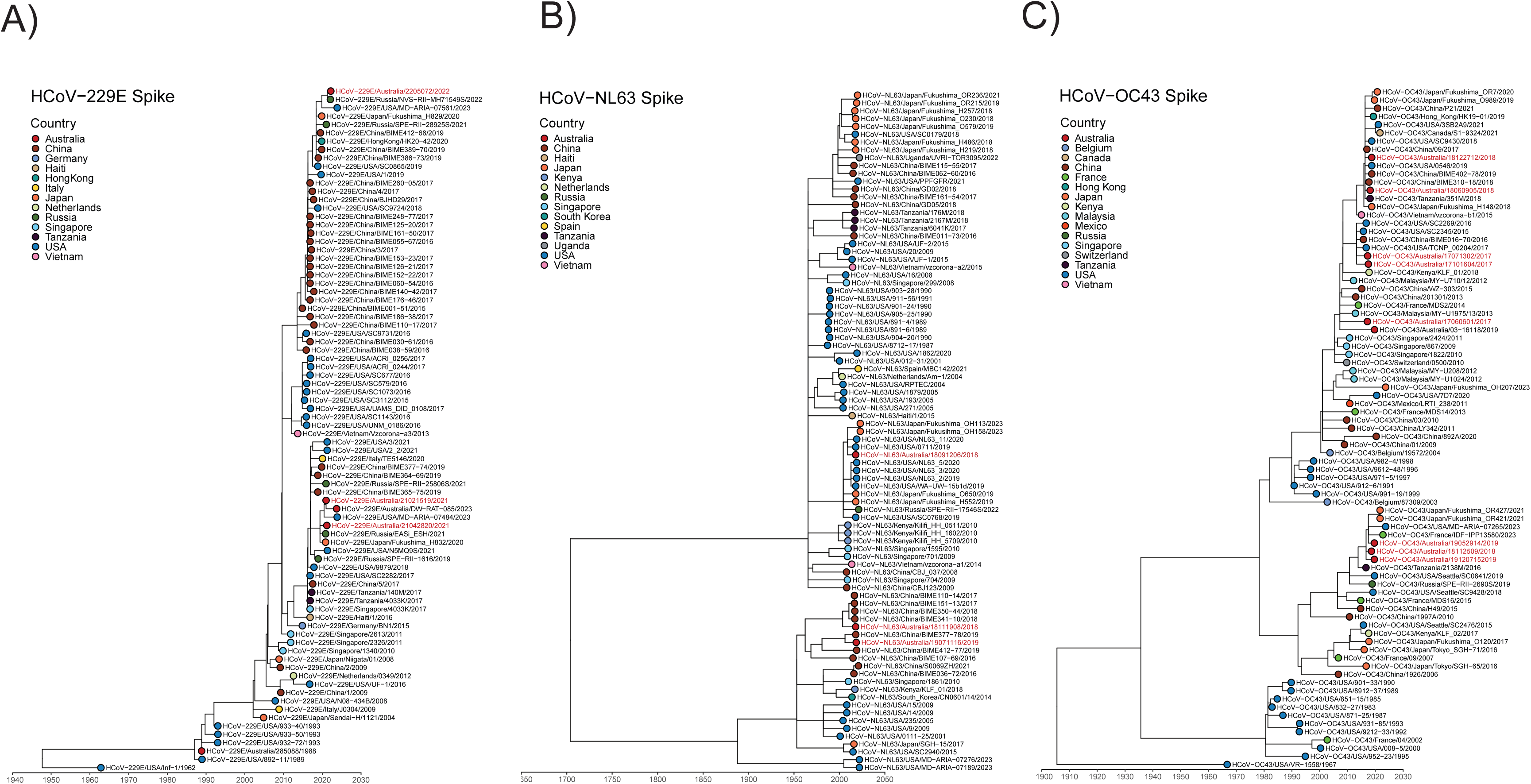
Alterations in sHCoV spike sequence. Time-resolved maximum likelihood phylogenetic analysis of spike nucleotide sequences of contemporary 229E (A), NL63 (B) and OC43 (C) isolates compared to publicly available sequences. Sequences from viruses isolated in this study are shown in red text. Coloured symbols indicate different countries of origin.

Analysis of NL63 S sequences showed 18111908, 19071116 and 18091206 grouped as they did in the full genome tree (Fig. 4B). Recombination analysis showed no evidence of recombination in our three NL63 isolates (data not shown). Analysis of OC43 S sequences showed all eight sequences grouped within the same genotype as the full genome tree (Fig. 4C). Recombination analysis of full genome OC43 sequences identified a recombination event in 17101604 between a genotype G and K sequence with breakpoints between S and Nucleocapsid (N) (Sup Fig. 3B).

### Growth of contemporary sHCoVs in immortalised cell lines

Downstream analysis of contemporary sHCoVs using supernatants from HNECs, BCi and AT2 cells is difficult as these culture systems produce small volumes of supernatant. Therefore, we attempted to establish protocols to produce larger quantities of virus using immortalised cell lines. For OC43, we attempted to grow the contemporary viruses in several cell lines including MRC-5, Huh7, HCT-8, BHK-21, BCi.NS1.1 and MvLu1 cells as well as Vero and A549 cells over-expressing human TMPRSS2. The lab-adapted OC43 strain caused CPE and had detectable virus growth (by qRT-PCR or antigen detection) in all cell types. However, virus growth was not observed for any contemporary OC43 isolates (data not shown). In contrast, the 229E isolates grew robustly and caused CPE in Huh7 cells and the NL63 isolates grew robustly and caused CPE in LLC-MK2 cells over-expressing ACE2 and TMPRSS2 (LLC-AT) as previously described (19, 20). Given the propensity of CoVs to mutate on passage in immortalised cell lines, we performed whole genome sequence analysis of 229E and NL63 isolates after two serial passages (P1 and P2) in Huh7 or LLC-AT cells, respectively. Comparison of NL63 isolate consensus sequences after two passages in LLC-AT cells showed the accumulation of two amino acid changes per virus isolate, one each in ORF1ab and one each in S (18091206: L1476F in ORF1ab and A408V in S, 18111908: H3489Y in ORF1ab and A428V in S and 19071116: L1945F in ORF1ab and P226L in S, Fig. 5A). Notably, all amino acid changes that occurred within S were in the NTD. In contrast, comparison of 229E isolate consensus sequences after two passages in Huh7 cells revealed no genetic and amino acid changes (Fig. 5B).

**Figure 5.**
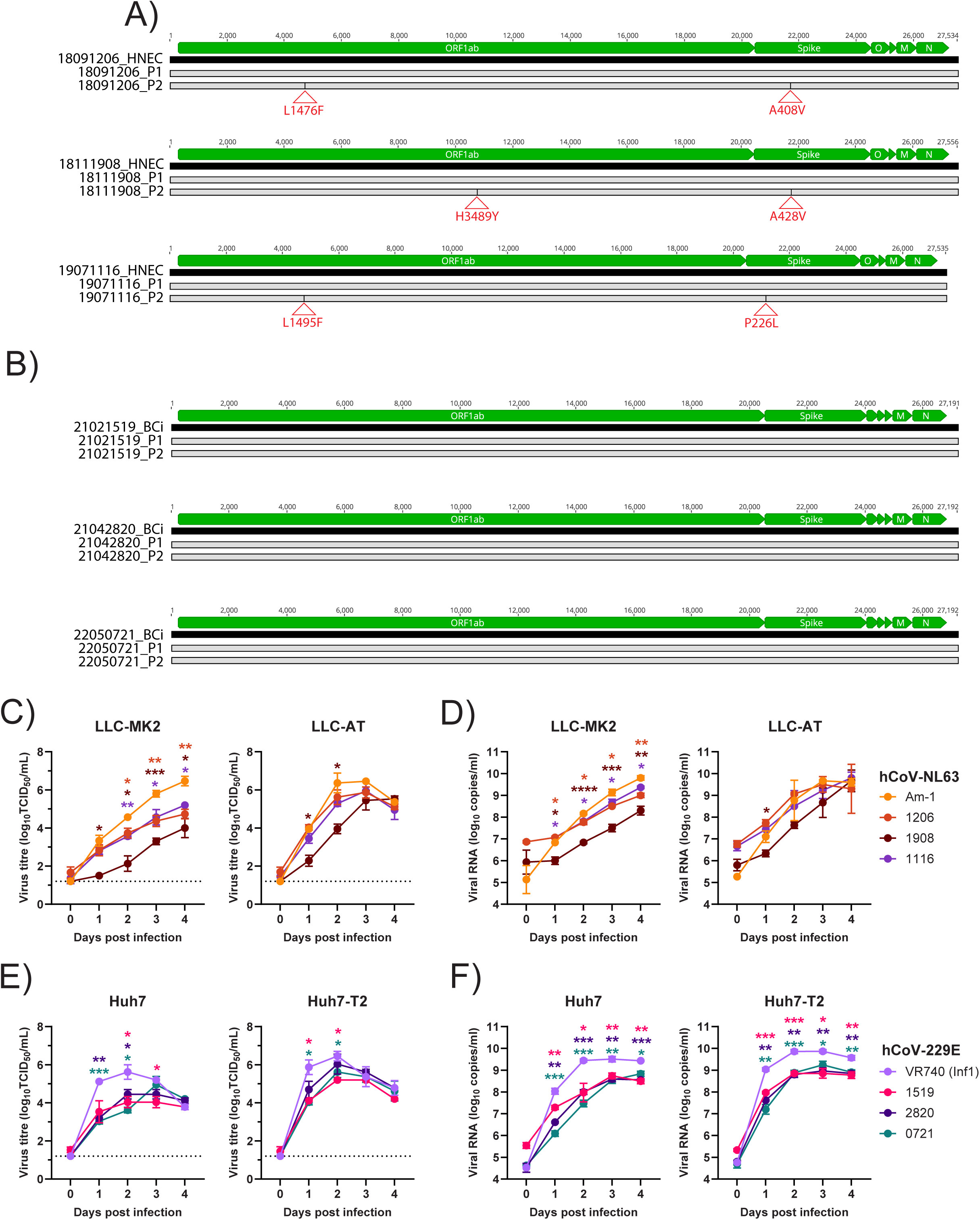
Comparison of virus growth in immortalised cell lines. Sequence alignments of A) NL63 and B) 229E virus stocks comparing passage 1 (P1) and passage 2 (P2) nucleotide sequences to the same virus isolated in HNEC (for NL63) and BCi (for 229E). Black lines indicate nucleotide changes, red triangles indicate amino acid changes. Infectious virus titres (C) and genome copies (D) in LLC-MK2 and LLC-AT cells infected with lab-adapted (Am-1) and contemporary NL63 isolates (18091206, 18111908, 19071116). Infectious virus titres (E) and genome copies (F) in Huh7 and Huh7-T2 cells infected with lab-adapted (VR740 (Inf1)) and contemporary 229E isolates (21021519, 21042820, 22050721). Data is a representative of at least 2 experimental repeats each with 3 replicates. Mean±SD is shown. Data were analysed using a two-way ANOVA with a Dunnett’s post-test. *p<0.05, **p<0.01, ***p<0.001 and ****p<0.0001.

To further explore the ability of contemporary sHCoVs to replicate in immortalised cell lines, we compared their replication kinetics (using P2 viruses) compared to the lab-adapted strains. All NL63 viruses replicated in LLC-MK2 cells with titres peaking at 4 dpi (Fig. 5C, D). However, Am-1 replicated to significantly higher titres than the contemporary isolates, with 18111908 having the lowest titres over the 4 days (Fig. 5C). Genome copies over time were consistent with infectious virus levels (Fig. 5D). Over-expression of both human TMPRSS2 and ACE2 in the cell lines accelerated growth of all NL63 viruses with infectious titres peaking at 2 dpi (Fig. 5C). While 18111908 still had slower growth compared to Am-1, the other two contemporary isolates grew to similar levels as Am-1 in both infectious virus titres and genome copies (Fig. 5C, D). Approximately 95-100% CPE was observed in LLC-AT cells by 3 dpi for all NL63 viruses. In comparison, CPE was only observed for Am-1 at 4 dpi in LLC-MK2 cells (data not shown). Overall, this suggests that the over-expression of ACE2 and TMPRSS2 significantly improves the replication of NL63 isolates.

All 229E viruses replicated in Huh7 cells with titres peaking at 2 dpi (Fig. 5E, F). Over-expression of human TMPRSS2 increased the titres of 229E at 1 dpi compared to Huh7 cells after which titres were similar between both cell types. Similar to the NL63 isolates, all contemporary 229E isolates replicated to lower titres compared to the reference strain VR740 in Huh7 cells. Over-expression of TMPRSS2 modestly increased the replication of 229E isolates so that their replication was similar in kinetics to VR740. Overall, this suggests that TMPRSS2 expression has a modest effect on 229E infection and Huh7 cells can be used to generate high titre stocks of 229E.

### Comparison of infection with lab-adapted vs contemporary isolates in the respiratory tract

We compared replication kinetics of lab-adapted and contemporary 229E and NL63 sHCoV strains across our human respiratory tract cell culture models. For NL63, both Am-1 and isolate 18111908 replicated similarly in HNECs as determined by genome copies but infectious titres were significantly lower for 18111908 compared to Am-1 over the 7 days (Fig. 6A, B). It is unclear whether this discrepancy is related to presence of defective genomes or ability to cause CPE in LLC-AT cells used in the infectivity assay. In contrast, only Am-1 replicated in BCi and AT2 cells (Fig. 6A, B). This is consistent with what we observed in isolating NL63 from nasal swabs -contemporary NL63 isolates preferentially infected the upper respiratory tract. In addition, it indicates a distinct difference in cell tropism between contemporary NL63 and lab-adapted NL63 viruses.

**Figure 6.**
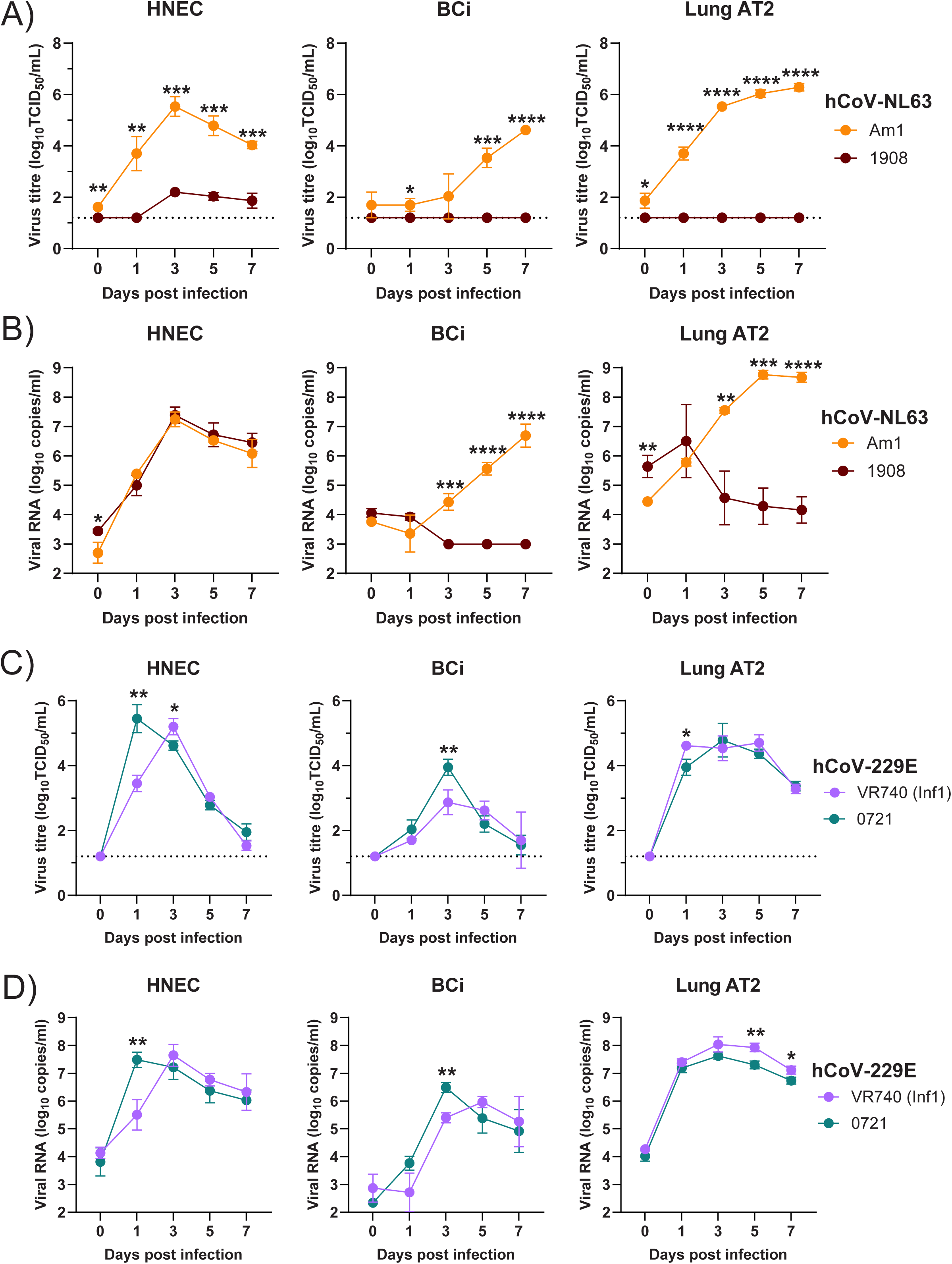
Comparison of virus growth across the human respiratory tract. Infectious virus titres (A) and genome copies (B) in HNECs, BCi and AT2 cells infected with lab-adapted NL63 (Am-1) or contemporary NL63 18111908 (1908). Infectious virus titres (C) and genome copies (D) in HNECs, BCi and AT2 cells infected with lab-adapted 229E (VR740-Inf1) or contemporary 229E 22050721 (0721). Data is a representative of 2 experimental repeats each with 3 replicates. Mean ± SD is shown. Data was analysed using a student’s t test. *p<0.05, **p<0.01, ***p<0.001 and ****p<0.0001

For 229E, both VR740 and 22050721 replicated in HNECs, BCi and AT2 cells (Fig. 6C, D). Clinical isolate 22050721 had higher titres at 1 dpi in HNECs and 3 dpi in BCi compared to VR740. In contrast, VR740 had modestly higher infectious virus titres on 1 dpi and genome copies on 5 and 7 dpi compared to 22050721 in AT2 cells. Again, this is consistent with what we observed in isolating 229E from nasal swabs - contemporary 229E viruses were able to infect all parts of the respiratory tract. We found that 22050721 shows similar replication kinetics as lab-adapted 229E.

To further understand how infection with contemporary sHCoVs differs from lab-adapted strains, we measured cytokine responses (IP-10, IFN-λ1, IFN-λ2/3 and IFN-β). In HNECs cells that were able to support replication of both lab-adapted and contemporary NL63, infection was associated with increased IP-10 secretion between 4 dpi and 6 dpi compared to the mock control, reaching significance only for Am-1 at 6 dpi (Fig. 7A). No significant difference in the level of IP-10 secretion was observed between strains and very little change in IFN-λ1, IFN-λ2/3 and IFN-β secretion was observed. Thus, NL63 appears to induce minimal induction of IP-10 and type I/III interferons, with a similar response between strains.

**Figure 7.**
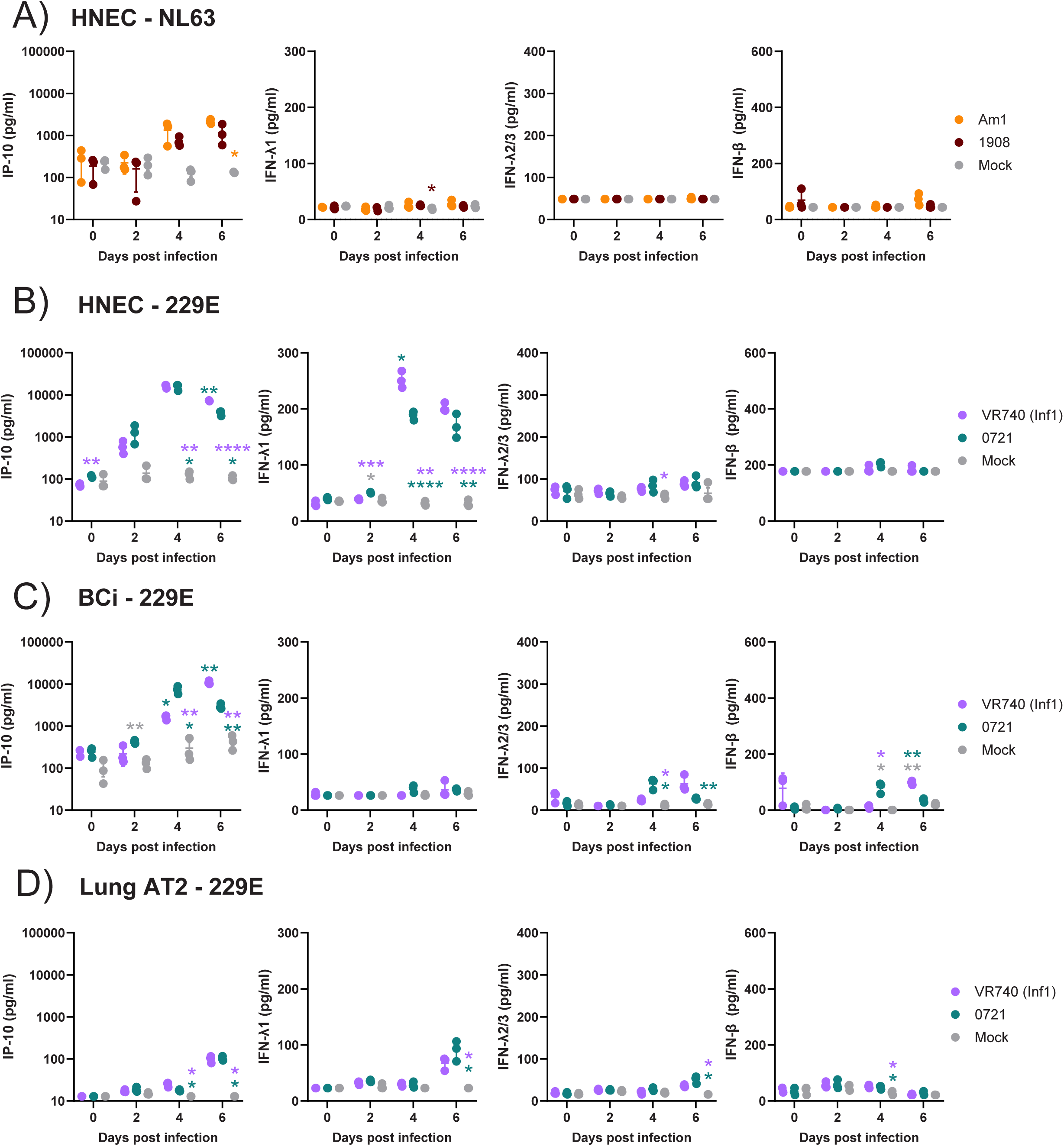
Comparison of the host response to sHCoVs. IP-10, IFN-λ1, IFN-λ2/3 and IFN-β1 levels in the supernatant of HNECs infected with NL63 18111908 or Am-1 (A), HNECs infected with 229E VR740 or 22050721 (B), BCi infected with 229E VR740 or 22050721 (C), and AT2 cells infected with 229E VR740 or 22050721 (D). Data is a representative of 2 independent experiments. Individual replicates and mean ± SD are shown. Data was analysed using a two-way ANOVA with Tukey’s post-test. *p<0.05, **p<0.01 and ***p<0.001.

In contrast to NL63, 229E infection strongly induced the secretion of both IP-10 and IFN-λ1 in HNECs compared to the mock control, peaking at 4 dpi (Fig. 7B). However, compared to VR740, IP-10 and IFN-λ1 secretion were lower on 6 dpi and 4 dpi, respectively, following 22050721 infection. As in the case of NL63, very little secretion of IFN-λ2/3 and IFN-β were observed in HNECs in response to 229E infection. In BCi, infection with 229E strongly induced IP-10 secretion peaking on 4 dpi for 2205298 and 6 dpi for VR740 (Fig. 7C). Modest increases in IFN-λ2/3 and IFN-β secretion were also observed with a similar peak for 2205298 on 4 dpi and VR740 on 6 dpi (Fig. 7C). In AT2 cells, modest increases in the secretion of all 4 cytokines were observed on 4 dpi and/or 6 dpi (Fig. 7D). However, no difference between cytokine secretion was observed between strains. Together these data indicate that infection with contemporary sHCoVs induces a similar cytokine profile following infection, but that kinetics and concentration can differ.

## Discussion

Here we demonstrated the utility of using specialised airway culture models to isolate novel contemporary sHCoVs isolates from human nasal swabs. Using these models we isolated 15 CoVs, from a total of 21 nasal swab specimens (71% success rate), which we further characterised for sequence variation, replication kinetics and host response compared to the commonly used lab-adapted reference strains. Previous work by Dijkman et al demonstrated the usefulness of air-liquid interface airway cultures for isolating sHCoVs. They recovered 9 viruses (4/6 HKU1, 4/4 OC43 and 1/2 229E) using primary bronchial epithelial cells (11). They also attempted to recover NL63 from 6 PCR+ nasal specimens but were only able to recover 1 isolate. This mirrors our observation that NL63 isolates are poorly recoverable in BCi cells. In another study, Komabayashi et al recovered 29 (16 OC43, 6 HKU1, 6 NL63 and 1 229E) CoVs from 36 nasal swab specimens in human bronchial epithelial cells and showed that OC43 and NL63 infected cultures produced infectious virus for at least 100 days (21). We did not measure viability nor propagate virus past 7 dpi. However, we did note that 229E caused obvious CPE in AT2 cells (data not shown) while no obvious CPE was observed with any other virus or cell type. In our study we did not have any samples that were PCR+ for HKU1 so we cannot comment on whether we could have recovered this virus using HNECs, BCi or AT2 cells.

Using our airway culture models, we noted that 229E isolates could be recovered or replicate in HNECs, BCi and AT2 cells, similar to the lab-adapted 229E strain. OC43 isolates were isolated in BCi which was in direct contrast to the lab-adapted strain that failed to replicate in these cells. NL63 isolates could only be recovered and replicate in HNECs whereas the lab-adapted strain could replicate in all airway cell types. Importantly, while HNECs require a nasal brush specimen from a healthy adult and generation of stem cell-derived AT2 cells can be challenging, our results suggest that the immortalized BCi-NS1.1 cell line differentiated at an ALI could be used to reliably recover 229E and OC43 isolates. Recovery of NL63 in HNECs is consistent with a higher expression of ACE2 in HNECs compared to BCi and AT2 cells (12, 22). This suggests that the tropism of the lab-adapted CoVs in the airways is distinct from those of clinical CoV isolates. Even for 229E that showed similar tropism between lab-adapted and contemporary CoV isolates, direct comparison revealed a slight difference in kinetics of viral replication and cytokine responses indicating the importance of incorporating contemporary sHCoVs in future research. A recent study by Otter et al demonstrated that replication kinetics, cellular tropism and cytotoxicity varied between 229E and NL63 lab-adapted strains and the pathogenic CoVs SARS-CoV-2 and MERS-CoV in HNECs (23). Given the differences we have observed with contemporary sHCoVs, there may be additional differences in replication kinetics, cellular tropism and cytotoxicity between pathogenic CoVs and the isolates described in our study.

While the ability to reliably recover CoV isolates is important, air-liquid interface cultures and stem-cell derived cultures are difficult and costly to scale up to produce virus stocks. Here we developed novel techniques to amplify and titrate virus stocks in immortalised cell lines for 229E using Huh7 cells and NL63 using LLC-AT cells. Importantly, we confirmed that there are very few amino acid changes observed for both 229E and NL63 after 2 passages in these cells indicating that virus stocks prepared in these cells can be used to reliably scale virus stocks for research (at least following 1-2 passages). The ability to measure the levels of infectious contemporary sHCoVs in infectivity assays will allow other researchers to study replication with these viruses. Komabayashi et al had also observed that NL63 isolates could be propagated in LLC-MK2 cells. However, we noted that while contemporary NL63 could also grow in LLC-MK2 cells in our hands, over-expression of ACE2 and TMPRSS2 increased viral titres and shortened the time to reach >80% CPE (4 days in LLC-AT vs >8 days for LLC-MK2) which was critical for measuring virus titres by infectivity assay.

Unfortunately, despite several attempts we could not develop an immortalised cell culture system to amplify and titrate contemporary OC43 isolates, though the lab-adapted strain replicates efficiently and causes CPE in many cell types. Other groups have also noted the difficulty of propagating OC43 clinical isolates in immortalised cell lines (24). It is notable that, like NL63 isolates, contemporary OC43 isolates show a clear difference in tropism within the respiratory tract compared to the lab-adapted reference OC43 strain. Whether this is related to differences in receptor usage or binding affinity, entry co-factor expression or replication machinery is unclear. However, our data suggest that studies investigating OC43 receptor usage should consider this discrepancy among isolates.

Of the three sHCoVs, NL63 showed the least genetic variability compared to the lab-adapted strain. This absence of genetic drift has been observed previously. The exclusivity of contemporary NL63 for replication in the nasal epithelium may shield NL63 from the selective pressure of antibodies in contrast to both OC43 and 229E contemporary isolates that can replicate beyond the upper respiratory tract. Overall, the greatest variability in sequence of the three sHCoVs compared to the lab-adapted reference strains was within the S gene. This is consistent with previous reports (8–10). For NL63, a single mutation within the S protein (I507L) has been associated with enhancing pseudovirus entry in cell culture (25). Whether the mutations we observed in ORF1ab and S following passage in LLC-AT cells are also associated with enhanced replication in cell lines requires further analysis.

Sequence analysis showed that most amino acid changes within our 229E isolates compared to the reference strain occurred within the RBD, particularly within the RBLs loops responsible for hAPN binding. Previous genetic and serological studies suggest substantial antigenic evolution has occurred within the 229E RBL over the last 60 years (8, 10, 26). Wong *et* al demonstrated that 229E RBDs sampled between the 1967 and 2015 could be classified into six classes based on RBL sequence (17). Interestingly, the RBD sequence of our isolates differed from RBDs categorised in these classes, suggesting further genetic evolution into a new RBD class. While Wong *et* al found that the more recent RBD classes (V and VI) bound hAPN with a higher affinity than classes I-IV, the affinity of more recent 229E RBDs (post-2015) requires further investigation. In contrast to 229E, NL63 isolates demonstrated minimal variation in the RBD compared to the reference strain. Instead, the Unique domain of NL63 showed the most variation in S1 compared to Am-1. While this domain is not required for binding to hACE2 (27), it has been speculated to bind to heparin sulfate as an attachment factor (28).

Recombination is an important mechanism of genetic variation in CoVs that can extend host range. Intraspecies recombination has been documented to occur in 229E, NL63 and OC43 sequences (10, 14, 15, 29). We identified one in three 229E isolates and one in eight OC43 isolates demonstrated evidence of genetic recombination. The 229E isolate (22050721) showed recombination with the co-circulating genotype 7b between ORF1ab (nucleotide 8152) and S (nucleotide 21534). Two studies found that 229E recombinants were rare, although several sequences were identified with breakpoints between S and M or N genes (30). Interestingly, recombination with breakpoints in ORF1ab and S have been demonstrated between HCoV-229E, bat 229E-related CoVs, and alpaca 229E-related CoVs (31). The OC43 isolate (17101604) showed evidence of recombination between S (nucleotide 24462) and N (nucleotide 29034). Recombination analysis of previous OC43 sequences by Pollett *et* al demonstrated S and N both had recombination breakpoints at similar positions to those 17101604 (32). While the number of isolates we analysed is small, we identified evidence of recombination in contemporary 229E and OC43 viruses.

Over-expression of TMPRSS2 increased the ability of all 229E isolates to replicate in Huh7 cells. For the contemporary isolates, over-expression of TMPRSS2 also meant that their replication was much more like that of the lab-adapted strain. A study by Shirato et al showed that two 229E clinical isolates, one from 2004 and one from 2008 were more dependent on TMPRSS2 for entry compared to the lab strain. They also showed that passage of a 229E clinical isolate 20 times through HeLa cells induced several mutations within the S gene that reduced TMPRSS2 usage and reduced replication fitness in human bronchial epithelial cells (33). A similar increase in replication was observed for the NL63 isolates in LLC-MK2 cells over expressing ACE2 and TMPRSS2. Others have shown that the early stages of NL63 entry into human airway epithelial cells, but not LLC-MK2 cells, is mediated by TMPRSS2 (34). Therefore, it’s unclear whether TMPRSS2 is contributing to the increased replication observed in LLC-AT cells or whether this increase is attributable to increased ACE2 expression.

There are some limitations to our study. Firstly, we did not attempt to recover virus from all nasopharyngeal swabs using all three respiratory models because we were constrained by the volume of sample. For example, we cannot definitively say that OC43 and 229E isolates can be recovered in HNECs since we only attempted to recover NL63 using these cells. However, given that we have shown 229E isolates replicate well in HNECs, we speculate that they could be recovered in HNECs. In addition, we did not attempt to recover all OC43 isolates in AT2 cells. Thus, while we failed to recover two, it is unclear whether these cells would be refractory to infection with all OC43 isolates. Our sample size for virus isolation experiments in primary HNECs was small (n=2 donors) and therefore we cannot speculate whether donor-dependent variation in the ability to recover isolates or replication efficiency of contemporary sHCoV isolates would occur. Finally, our experiments examining replication kinetics and cytokine expression in the respiratory tract only compared one contemporary isolate each to the lab-adapted reference strain. Given the genetic diversity between the isolates, particularly for 229E, it is possible that the isolate selected for analysis may not reflect all circulating 229E isolates.

Overall, here we have shown that contemporary sHCoVs differ from lab-adapted reference strains and should be used for study of virus biology and evaluation of medical countermeasures. Furthermore, the technical advances described in our study should make working with these HCoVs easier and facilitate future research endeavours.

## Materials and Methods

### Cell Lines

MRC-5 cells (CCL-21, ATCC) were maintained in DMEM supplemented with 50 U/mL penicillin, 50 µg/mL streptomycin, 1 mM sodium pyruvate, 20 mM HEPES, 0.1 mM non-essential amino acids, 2 mM Glutamax, 0.18% (v/v) sodium bicarbonate and 10 % (v/v) FBS. Huh7 cells were maintained in DMEM supplemented with 50 U/mL penicillin, 50 µg/mL streptomycin, 2 mM Glutamax and 10 % (v/v) FBS. LLC-MK2 cells were maintained in MEM supplemented with 50 U/mL penicillin, 50 µg/mL streptomycin, 15 mM HEPES, 2 mM Glutamax, and 5% (v/v) FBS. Basal-like human airway progenitor cells (BCi-NS1.1, (35)) were maintained in Pneumacult-Ex Plus medium (StemCell Technologies, Cat. 05040) supplemented with 96 ng/mL Hydrocortisone (StemCell Technologies, Cat. 07925) and 50 U/ml Penicillin and 50 µg/ml Streptomycin.

### Genetically modified cell lines

LLC-AT cells were generated by transducing LLC-MK2 cells with lentiviral constructs expressing human ACE2 (pHAGE2 containing the angiotensin-converting enzyme 2 gene; NR-52512, BEI Resources, NIAID, NIH) and TMPRSS2 (pscALPSblasti-TMPRSS2 Blasti, Addgene plasmid #158088, a gift from Jeremy Luban). LLC-AT cells were maintained in MEM supplemented with 50 U/mL penicillin, 50 µg/mL streptomycin, 15 mM HEPES, 2 mM Glutamax, and 5% (v/v) FBS. Huh7 overexpressing TMPRSS2 (Huh7-T2) were made by transducing Huh7 cells with a lentiviral construct expressing TMPRSS2 (pscALPSblasti-TMPRSS2 Blasti, Addgene plasmid, Cat. 158088, a gift from Jeremy Luban). Transduced cells were selected by supplementation of culture media with 2 µg/ml Blasticidin (Gibco, Cat. A1113903) until negative control cells were dead. TMPRSS2 expression was confirmed by surface staining selected cells with 5 µg/ml goat anti-TMPRSS2 PE (BioLegend, Cat. 378403) antibodies and analysed by flow cytometry. Huh7-T2 cells were maintained in DMEM supplemented with 50 U/mL penicillin, 50 µg/mL streptomycin, 2 mM Glutamax, 10 % (v/v) FBS and 1 µg/ml Blasticidin.

### Nasal brushing for nasal epithelial cells

Healthy adults were recruited for this study under ethics approval HREC/35132. Nasal epithelial cells were sampled and cultured in Pneumacult-Ex Plus medium (StemCell Technologies, Cat. 05040) supplemented with 96 ng/mL Hydrocortisone (StemCell Technologies, Cat. 07925), 50 U/ml Penicillin and 50 µg/ml Streptomycin and 0.25 µg/ml Amphotericin B (ThermoFisher Scientific, Cat. 15290018) as described previously (12, 36, 37).

### Differentiation of airway epithelial cells

Nasal epithelial cells or BCi-NS1.1 cells were seeded at 300,000 cells/well in 12mm transwell membrane supports (Corning, Cat: 3460) or 150,000 cells/well in 6.5mm transwell membrane supports (Corning, Cat: 3450) and cultured in Pneumacult-Ex Plus medium in apical and basolateral chambers. Upon reaching confluence (2-3 days post-seeding), apical medium was removed to expose cells to ambient air; medium in the basolateral chamber was replaced with Pneumacult ALI maintenance medium (Stemcell Technologies, Cat: 05001) supplemented with 4 µg/mL Heparin (Stemcell Technologies, Cat: 07980), 480 ng/mL Hydrocortisone and 50 U/ml Penicillin and 50 µg/ml Streptomycin. HNEC cultures were also supplemented with 0.25 µg/mL Amphotericin B. Medium in the basolateral chamber was replaced with fresh Pneumacult ALI maintenance medium every 2 to 3 days. The apical surfaces of cultures were washed with PBS (+Ca^2+^/Mg^2+^) once a week to remove mucus accumulation. BCi cells were cultured for a minimum of 28 days and to form differentiated, polarized human bronchial epithelial cell cultures, while nasal epithelial cells were cultured at an ALI for 42 days to form differentiated nasal epithelial cell (HNEC) cultures. Amphotericin B was removed from basolateral medium at 28 days post-airlift.

### Generation of hESC-derived AT2 cells

Human embryonic H9 (female) stem cells were differentiated into lung AT2 cells as previously described (12, 38). Briefly, H9 stem cells were differentiated as a three-dimensional organoid for 18 days in flasks coated with Matrigel (Corning, Cat: 354230). Lung organoids were maintained for experiments between passage 2-8 and dissociated with TrypLE (Thermo Fisher Scientific, Cat: 12604013) and seeded onto Geltrex-coated plates (Gibco, Cat: A1413201). Cells were then maintained for a further 7-10 days in two-dimensional culture until infection at >70% confluency.

### Lab virus Stocks

NL63 Lab virus (Amsterdam-1) was a kind gift from Prof Lia van der Hoek, University of Amsterdam, shared by Prof Kirsten Spann, Queensland University of Technology with permission. Viral stocks were prepared from supernatants of infected LLC-MK2 cells. 229E (VR740 (Inf1)) and OC43 (VR1558) were kind gifts from Prof Nathan Bartlett, University of Newcastle. Viral stocks were prepared from supernatants of infected MRC-5 cells.

### Clinical Material

Stored nasopharyngeal specimens were shared by the Victorian Infectious Disease Reference Laboratory, Melbourne, Australia for the isolation of contemporary HCoVs as approved by the University of Melbourne Human Ethics Committee (Reference 2024-31337-61425-3). Briefly, samples were collected by referring pathology services and stored at –80°C. Samples were typed using the following primers CV229e-F 5’ TCACATGTTGTACGGCTAGTGATAAA 3’, CV229e-R 5’ ACCCACCATTTGAATAAACAACCT 3’, probe CV229e 5’ VIC-AGCAAGCTCATTACTAAGTCTA-MGBNFQ 3’, HCor-F 5’ AAATTTTATGGTGGCTGGAATAATATGTT 3’, HCoronaRT-Ra 5’ TAGGCATAGCTCTRTCACAYTT 3’, HCoronaRT-Rb 5’ TTGGCATRGCACGATCACAYTT 3’ and probe Corona 5’ FAM-TGGGTTGGGATTATC-MGBNFQ 3’ which can detect the presence of 229E or other non-229E, non-HKU1 CoVs (either NL63 or OC43). No samples containing HKU1 were analysed. A total of 21 samples were used to recover infectious HCoVs: 5 229E and 16 other HCoVs. Two samples were lost due to fungal contamination. A list of the samples and their collection dates is shown in Table 1.

### Isolation of infectious virus from clinical specimens

200 µL of clinical specimen was diluted with 600 µL of PBS and centrifuged at 2000 rpm for 5 minutes. Clarified sample (200 µL) was added to each well of either airway epithelial cells or AT2 cells and incubated for 2 h at 33°C. Following virus adsorption, the virus inoculum was removed, and cells were incubated at 33°C for up to 7 days. AT2 cells were cultured in 500ul of media. Supernatant was collected daily and 500 µL of media re-added to the cells. For airway epithelial cells, 200 µL of PBS was added to the apical surface, incubated for 10 min at 33°C and collected. All viruses were stored at -80°C.

### Next-generation sequencing

Stocks of NL63, 229E and OC43 were extracted using the QIAamp Viral RNA Mini Kit. Complementary DNA (cDNA) was synthesised from 15μl of each sample using random hexamers and the ProtoScript II first strand cDNA synthesis kit (New England Biolabs, Ipswich, MA, USA), followed by the NEBNext Ultra II Non-Directional RNA Second Strand Synthesis Module (New England Biolabs). Indexed libraries were prepared using the Twist Total Nucleic Acids Library Preparation Kit for Viral Pathogen Detection and Characterization (Twist Biosciences, South San Fransisco, CA, USA) following the manufacturer’s protocol. Pre-capture libraries were quantified using QubitTM dsDNA HS (Thermo Fisher Scientific) and qualified using Agilent Tapestation DNA HS reagents (Agilent Technologies, Santa Clara, CA, USA). Indexed libraries were pooled by mass prior to capture. Twist hybridisations followed the Twist Target Enrichment Standard Hybridization v1 Protocol using the Twist Comprehensive Viral Research Panel, according to manufacturer’s instructions and with a 16-hour incubation at 70°C. Libraries were quantified and qualified as above, before sequencing using a paired-end 150 bp chemistry with a P1 cartridge on an Illumina NextSeq2000 (Illumina, San Diego, CA, USA).

### Sequence analysis

Reads were mapped to viral reference genomes for relevant taxa accessed from NCBI Virus (https://www.ncbi.nlm.nih.gov/labs/virus/vssi/#/) iteratively using Minimap2 to generate consensus sequences (39). To generate sequence similarity plots (Simplots), full genome nucleotide sequence alignments were input into Simplot++ for Simplot analysis using the Jukes Cantor substitution model with a window size of 100bp and a step size of 20 bp (40).

### Phylogenetic analysis

Sequences were aligned using MAFFT version 7.526 (41). Time-resolved trees were constructed using maximum likelihood method with generalized-time reversible (GTR) nucleotide substitution model as implemented in IQ-TREE2 version 2.3.3 (42) and least-square dating method (43). Trees were visualised using ggtree version 3.10.1 (44) and sequences were grouped by lineage or genotype as described by Ye at al. (14).

### Recombination analysis

Sequence alignments generated using MAFFT version 7.526 (41) were analysed using recombination detection program version 5 (RDP5) (45). Similarity plots were generated using the RDP Method and 30 bp window size.

### Production of 229E and NL63 viral stocks in immortalised cell lines

Airway epithelial cell supernatants (HNEC-derived NL63 and BCi-derived 229E) were diluted with MEM (1:10) and inoculated onto confluent flasks of either LLC-AT or Huh7 cells for 1 h at 33°C. Following adsorption, additional media (1% FBS and 1 μg/mL TPCK-trypsin for Huh7 and 2% FBS and 1 μg/mL TPCK-trypsin for LLC-AT) was added. Flasks were cultured until approximately 80-100% CPE was observed (day 3-4 for 229E and day 6-7 for NL63). Supernatants were clarified by centrifugation at 4000 rpm for 10 mins and virus stock aliquots stored at -80°C. Stock titres were determined by virus titration as described below.

### Replication kinetics in immortalised cell lines

Confluent monolayers of Huh7, Huh7-T2, LLC-MK2 and LLC-AT cells seeded into 24 well plates were washed with 500 µL plain MEM before inoculation with 100 µL of either 229E diluted to a multiplicity of infection (MOI) of 0.001 or NL63 diluted to an MOI of 0.01 for 1 hour at 33°C. Following removal of the virus inoculum, the cells were washed 3 times and cultured at 33°C up to 4 dpi. Supernatants were collected daily for virus titration and qRT-PCR as described below.

### Virus infection of human airway epithelial cells

Prior to infection, airway cultures were washed once with pre-warmed PBS (+Ca^2+^/Mg^2+^) to remove mucus on the apical surface. Cells were inoculated with 100 µL 229E diluted to an MOI of 0.01, NL63 diluted to 10^7^ genome copies, or OC43 diluted to an MOI of 0.1 for 2 hours at 35°C. The inoculum was removed, and the apical surface washed three times with PBS and the third wash collected as the day 0 sample. On days 1-7 post-infection, 200 µL PBS was added to the apical chamber and incubated at 35°C for 10 minutes before samples were collected and stored at -80°C. Basolateral medium was replaced every second day with ALI maintenance medium. Apical samples were assayed by virus titration as described above and qPCR (E gene copies) as described below.

### Virus infections of lung AT2 cells

For infection of lung AT2 cells, cells were inoculated with 10^4^ TCID_50_ of 229E, 10^7^ genome copies of NL63 or 10^4^ TCID_50_ OC43 in 100 µL for 1 hour at 35°C. Cells were washed three times with PBS, the third wash was collected as the day 0 sample and cells were refed with lung AT2 medium. Supernatant samples were collected every day for seven days and the media was replenished on days 1-6. Supernatant samples were assayed by virus titration and qRT-PCR as described below.

### Quantification of infectious virus titre

Virus titrations were performed in 96-well plates with confluent monolayers of LLC-AT, Huh7 cells and Huh7-T2 cells. For Huh7 and Huh7-T2 cells, plain DMEM was used to wash cells and replaced with 180 μL media containing 1% FBS and 1 μg/mL TPCK-trypsin. For LLC-AT cells, plain MEM was used to wash cells and replaced with 180 μL media containing 2% FBS and 1 μg/mL TPCK-trypsin. Each sample was titrated in quadruplicate by adding 20 μL supernatant to the first well and performing 10-fold serial dilutions. Cells were incubated at 33°C and assessed microscopically for virus-induced cytopathic effect (CPE) on day 3 for 229E or day 7 for NL63. Virus titers are expressed as mean log_10_TCID_50_/mL.

### RNA Extraction and qRT-PCR

RNA was extracted as per the manufacturer’s recommendation using either the QiaCube HT (Qiagen) and QiaAmp 96 Virus QiaCube HT kit (Qiagen, Cat. 57731) or QIAamp Viral RNA Mini Kit (Qiagen, Cat. 52904). qRT-PCR reaction was set up using SensiFast Probe No-ROX One-Step Kit (Bioline, Cat. BIO-76005) using the following primers/probes: NL63 Forward: 5’-AGGTTGACTTGTATAATGGTGCT-3’, NL63 Reverse: 5’-GCCAACACAAAGAAAAATATCA-3’ and NL63 Probe: 5’-TGCCGAAGAGCCTGTTGTTGGT -3’; 229E Forward: 5’-ATGTGTACCACATTTACCAATCA-3’, 229E Reverse: 5’- TCCAAACTGAAGAATAACAATGA -3’ and 229E Probe: 5’-TGCACATAGACCCTTTCCCTAAACG-3’; OC43 Forward: 5’-TGTTTATGGCTGATGCTTATCT -3’, OC43 Reverse: 5’-AAGGTATTACACATACCGCAAA -3’ and OC43 Probe: 5’-CACTGTGTGGTATGTGGGGCAAAT-3’. Serial 10-fold dilutions of plasmid encoding the viral sequence target of interest were used to generate a standard curve and calculate the virus genome copies in the samples.

### Cytokine bead array

Cytokine/chemokine concentrations in supernatants or basolateral medium were analysed using the LEGENDplex human anti-virus response panel (Biolegend, Cat. 740390) following the manufacturer’s instructions. Samples were run on a BD FACSCanto II and analysed using LEGENDplex™ Data Analysis Software Suite.

## Acknowledgements

This work was supported by a foundation grant from the Cumming Global Centre for Pandemic Therapeutics (CGCPT00021). K.S. was supported by a National Health and Medical Research Council of Australia Investigator grant (APP1177174). We thank Dr Ronald Dijkman for helpful discussions regarding isolation of sHCoVs from respiratory specimens. We are grateful to Drs Nathan Bartlett, Kirsten Spann and Lia van der Hoek for sharing the reference CoVs. We thank the Victorian Infectious Disease Reference Laboratory for their assistance with the next-generation sequencing of sHCoVs.

## Figures Legends

**Sup. Figure 1. Isolation of contemporary sHCoVs.** Log_10_ genome copies/mL of NL63-18091206 (A) and OC43 isolates in AT2 cells (B) demonstrating lack of virus recovery. Log_10_ genome copies/mL of lab-adapted OC43 (VR-1558) in BCi (C) and AT2 cells (D) from 0 to 7 dpi. For (A), the mean from 4 pooled replicates is shown. For (B), (C) and (D), the mean ± SD from 4 un-pooled replicates is shown. (A) and (B) represents data generated from a single attempt to recover virus using a nasal swab specimen. (C) and (D) are representatives of two independent experiments each with 3 replicates.

**Sup. Figure 2. Sequence analysis of sHCoVs.** Analysis of 229E S1 (D), NL63 S1 (E) and OC43 S1 (F) from contemporary isolates compared to the reference strain. Protein domains are annotated in blue and receptor binding loops (RBLs) or sialic acid binding loops (SBL) are annotated in orange. Dots indicate the same amino acid as the reference sequence.

**Sup. Figure 3. Recombination analysis of sHCoVs.** Recombination pattern and breakpoints of A) HCoV-229E/Australia/22050721/2022 with China/BIME365-75/2019 and Japan/Fukushima_H829/2020, and B) HCoV-OC43/Australia/17101604/2017 with Kenya/KLF_01/2018 and Japan/Fukushima_H148/2018. The recombinant region is shown in red with the 95% and 99% breakpoint confidence intervals shown in dark and light grey, respectively.

