## Supplemental Figures for "Contemporary seasonal human coronaviruses display differences in cellular tropism compared to laboratory-adapted reference strains"

### Supplementary Figure 1 - Isolation of contemporary seasonal hCoVs

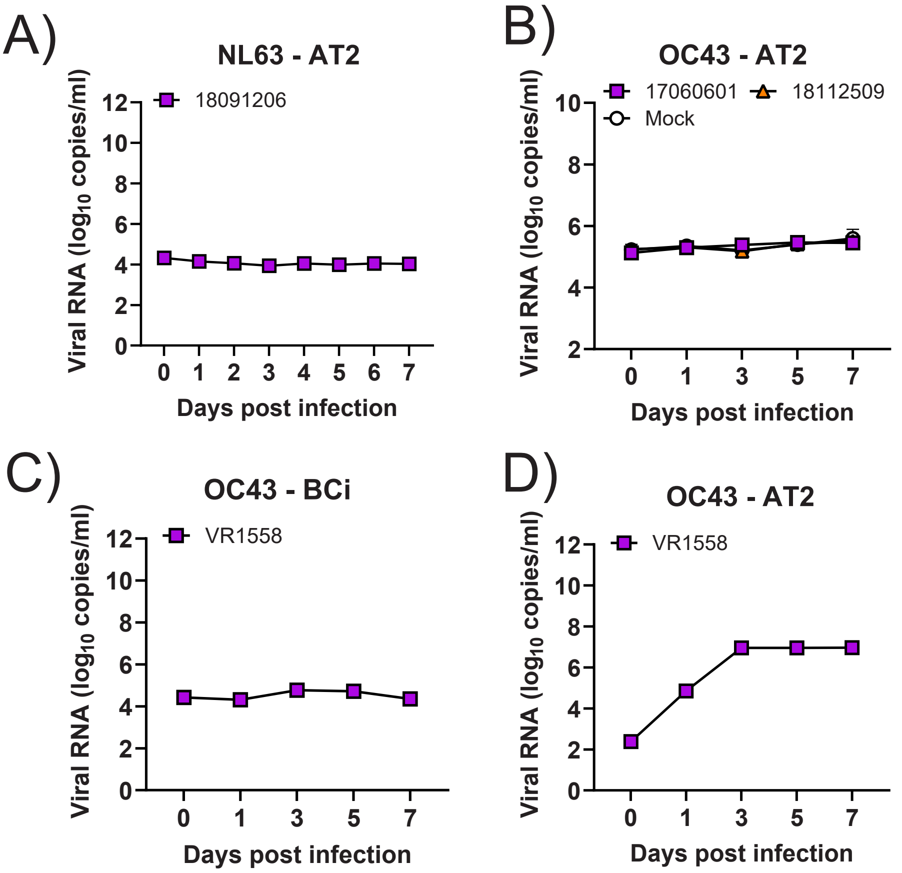

### Supplementary Figure 2 - Alignment of S1 sequences

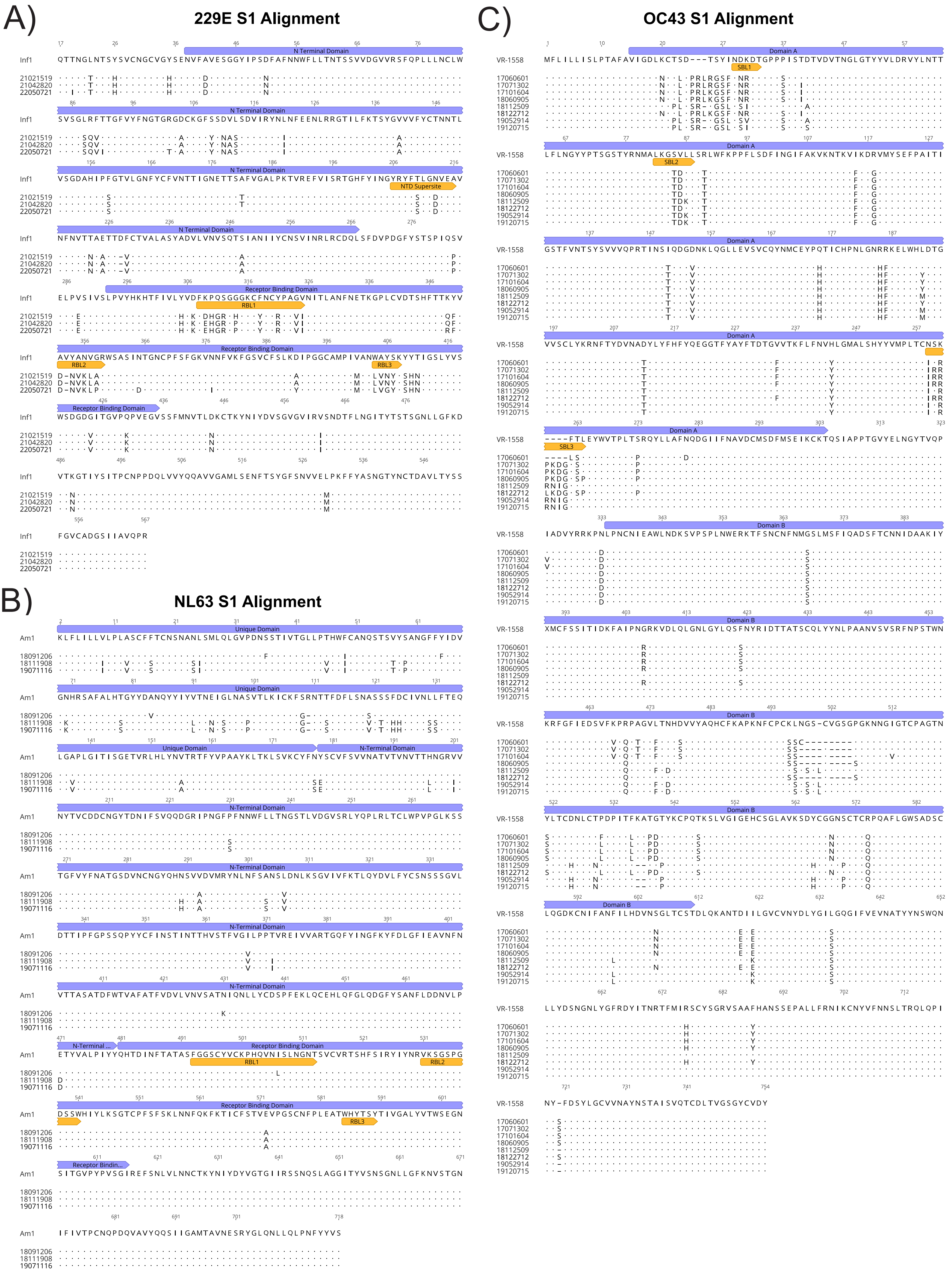

### Supplementary Figure 3 - Recombination Analysis

A)

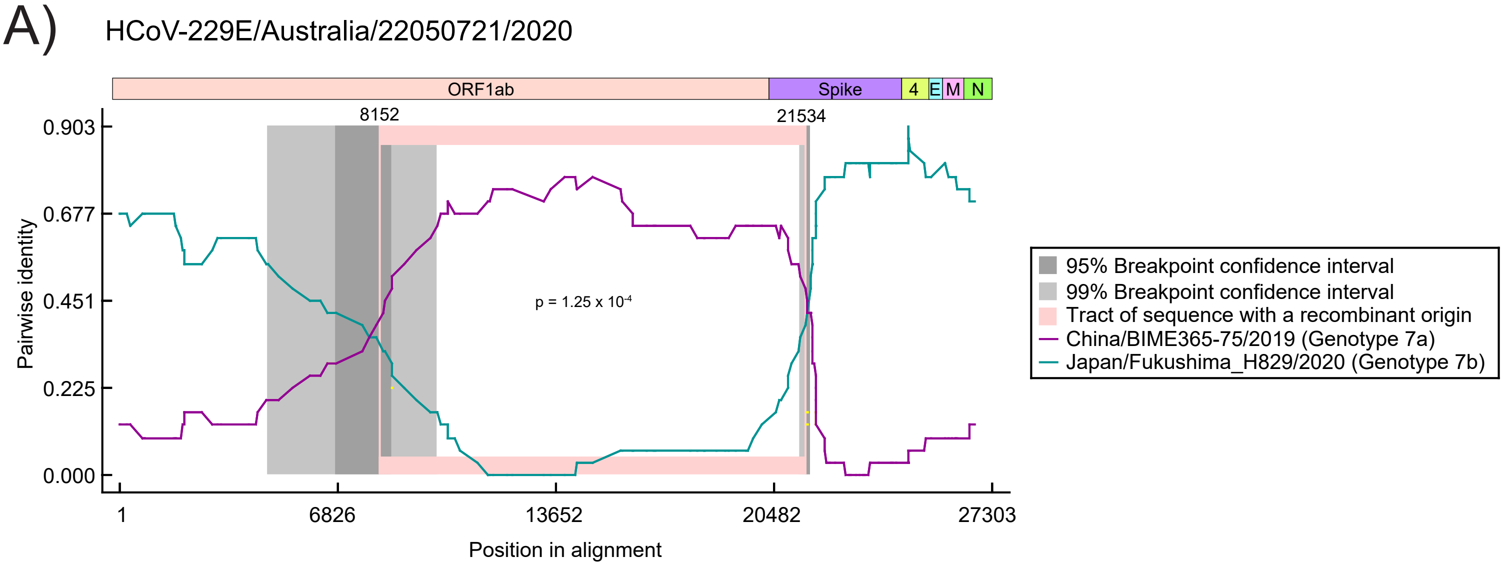


B)

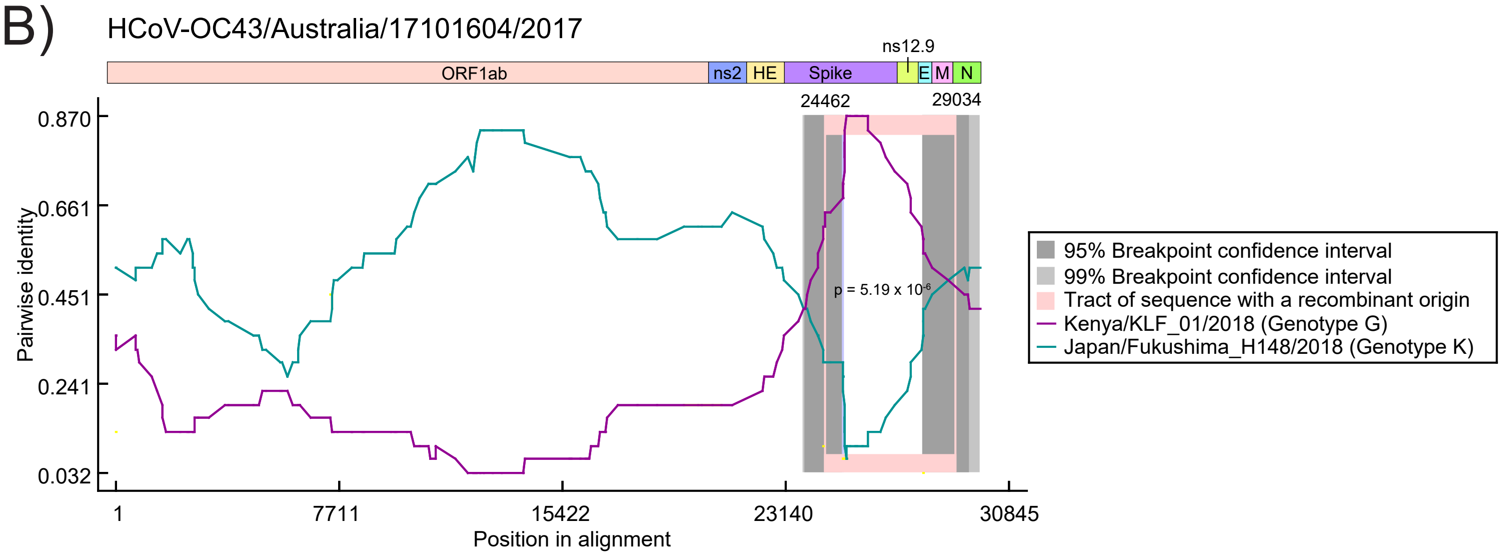
